# Defective cell death of distinct microglial subsets contributes to ADHD-like behavior in mice

**DOI:** 10.1101/749390

**Authors:** Hsiu-Chun Chuang, Eva K. Nichols, Isabella Rauch, Wei-Cheng Chang, Rhea Misra, Patrick M. Lin, Maiko Kitaoka, Russell E. Vance, Kaoru Saijo

## Abstract

Microglia are resident immune cells in the central nervous system that play essential roles to maintain homeostasis and neuronal function. Microglia are heterogeneous cells but the mechanisms by which they contribute to normal brain development remain unclear. Here,we show that microglia in the developing striatum and thalamus undergo pyroptosis,a type of lytic cell death that occurs as a result of Caspase-1 (CASP1) activation downstream of inflammasomes. We observe that pyroptosis occurs in a spatiotemporally regulated and Casp1-dependent manner during fetal brain development. Mice lacking *Casp1* or the inflammasome regulating molecules, NLRP3, IL-1R, and Gasdermin D exhibit behavior changes characterized by hyperactivity, inattention, and impulsivity that are similar to attention-deficit/hyperactivity disorder (ADHD). Furthermore, re-expression of *Casp1* in Cx3cr1^+^ cells including microglia restores normal behavior and cell death. We demonstrate that injection of an NLRP3 inhibitor into pregnant wild-type mice is sufficient to induce ADHD-like behaviors in offspring. These data suggest that microglial inflammasome activation and pyroptosis are essential for normal brain development and that genetic and pharmacological disruptions in this pathway may represent new ADHD risk factors.

---

Microglia cells serve as sentinels to manage central nervous system (CNS) injury and infection while maintaining homeostasis by influencing neuronal development, viability, and function. Recent single-cell RNA-sequencing analyses demonstrate that microglia in the developing brain are highly heterogeneous^1-3^ suggesting that specific microglial subsets may have distinctive functions for brain development, and that deregulation of microglial subsets could lead to discrete neurodevelopmental disorders.

Pyroptosis is a lytic cell death that occurs downstream of inflammasomes, which are cytosolic sensors for a variety of pathogenic and noxious stimuli^4^. Activated inflammasomes initiate the dimerization and cleavage of pro-CASP1 to active CASP1, which then cleaves pro-interleukins (IL)-1β and pro-IL-18 to generate mature cytokines. CASP1 also cleaves and activates Gasdermin D (GSDMD), which in turn promotes pyroptosis, IL-1β secretion, and inflammation^5,6^ (Sup. Fig. 1a). Other types of cell death such as apoptosis plays essential roles in the early development of several organs including the brain^7^, yet, whether or not pyroptosis is required for brain development remains uncertain. Interestingly, mutations in the inflammasome protein *NLRP1* and IL-1 receptor 2 (IL-1R2) are found in patients with autism spectrum disorders (ASD)^8,9^ and IL-1 receptor 1 (IL-1R1) has been implicated in hyperactivity in mice^10^. Therefore, we hypothesized that microglial pyroptosis pathways may be activated during normal brain development, and that alterations in the pathway could be involved in behavior changes such as ASD or hyperactivity.

To investigate microglial pyroptosis during fetal brain development, we developed an assay to image microglial death in fetal brains. We used propidium iodide (PI) to visualize cells undergo lytic cell death, such as pyroptosis^11^. PI was injected into pregnant WT females crossed with *Cx3cr1*^GFP/GFP^ male mice at specific days during fetal brain development. Genotype of offspring is *Cx3cr1*^GFP/+^ in which microglia are labeled by GFP^12^. While it is known that *Cx3cr1* is expressed in non-microglial cells outside the brain, *Cx3cr1*-GFP^+^ cells in the brain parenchyma are considered microglia^13^. Embryos were recovered, cleared using the CUBIC method^14^, and their brains were imaged via light sheet fluorescence microscopy (LSFM; Fig. 1a). Interestingly, our imaging revealed discrete clusters of PI^+^GFP^+^ cells in both the lateral ganglionic eminence (LGE), which matures into a major part of striatum, and the intermediate thalamus (iTh) starting at E12.5 (Fig. 1b and Sup. Fig. 1b-d). These clusters were also noted in both the LGE and thalamus at E14.5 (Fig. 1c, e, and Sup. Movie 1), coincident with active neurogenesis and neural cell death^15^, and were almost absent at E16.5 (data not shown). To investigate whether these clusters were indeed associated with inflammasome formation and thus pyroptosis, we stained fetal brain tissue with an anti-ASC (apoptosis-associated speck protein containing a CARD) antibody^16^ and observed ASC specks that are characteristic of inflammasome formation in the clusters (Sup. Fig. 1e-h). Since Casp1 activation is required for pyroptosis (Sup. Fig. 1a), we next tested whether cell death is impaired in *Casp1*^*-/-*^ mice. Using *Cx3cr1*^GFP/GFP^ mice crossed with *Casp1*^*-/-*^ mice, we found that formation of PI^+^GFP^+^ clusters in the LGE and iTh were significantly reduced in *Casp1*^*-/-*^; *Cx3cr1*-GFP offspring (Fig. 1d, e, and Sup. Movie 2). These data suggest that microglia in the LGE and iTh, undergo pyroptosis during fetal brain development.

**Fig. 1.**
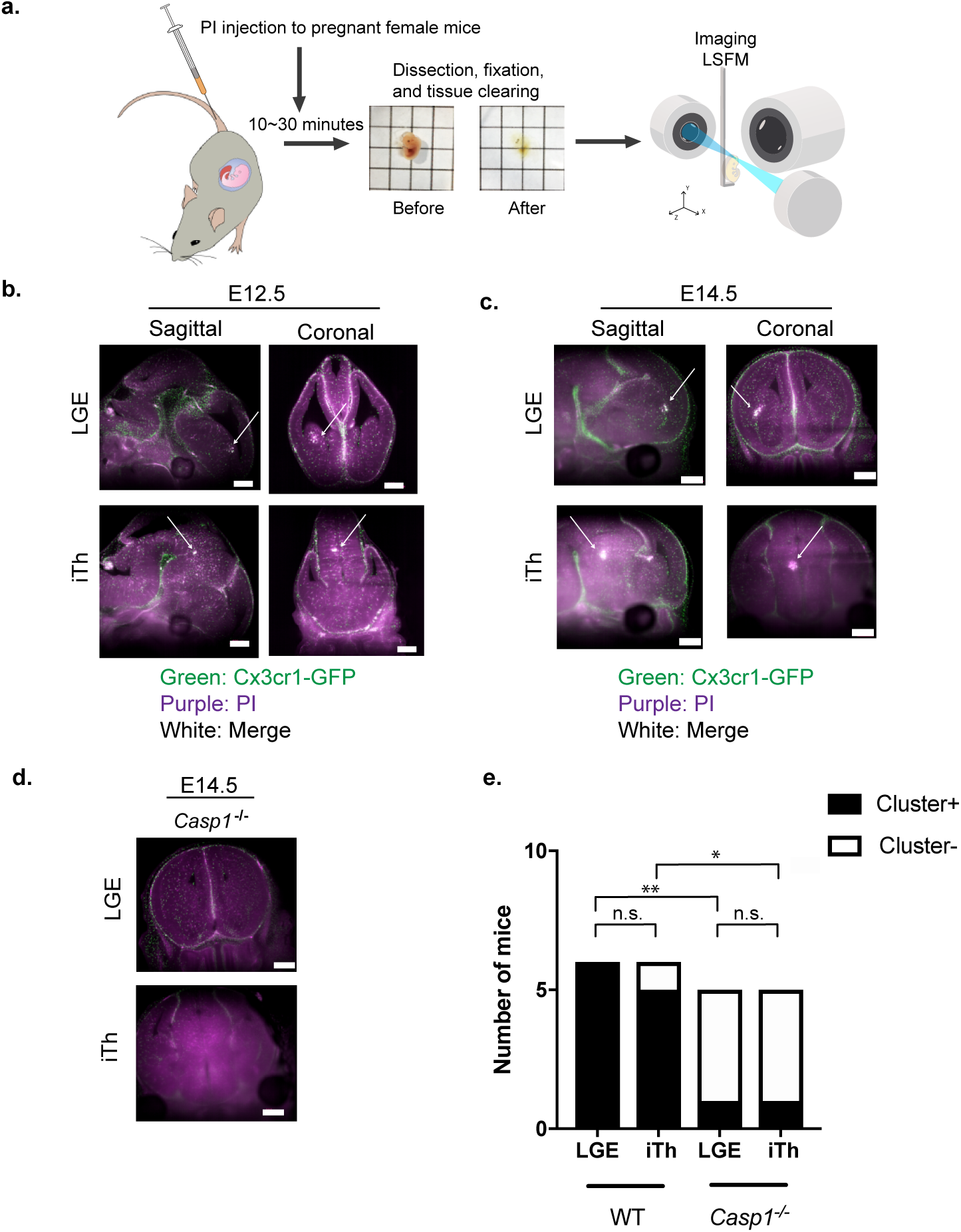
Microglial pyroptosis occurs in a spatiotemporal manner in the fetal brain. **a.**Experimental scheme is shown. Propidium Iodide (PI) was injected into pregnant female mice at specific gestational stages. Ten to thirty minutes after PI injection, total embryos were recovered, cleared, and imaged by LSFM. **b.** Representative sagittal and coronal LSFM images of the LGE and primitive thalamus (iTh) of an E12.5 fetal brain are shown (N=6). Arrows indicate the location PI+ cluster. Green: GFP, Purple: PI, White: Merge. Scale bars indicate 500 μm. **c.** Representative images of an E14.5 fetal brain are shown (N=6). Arrows indicate the location of PI+ cluster. Green: GFP, Purple: PI, White: Merge. Scale bars indicate 500 μm. **d.** Representative coronal LSFM images of *Casp1*^*-/-*^ at E14.5 are shown (N=5). Scale bars indicate 500 μm. **e**. Numbers of mice that exhibited clusters in the LGE and iTh are shown. Asterisk (*) shows p<0.05 and (* *) shows p<0.01.

In order to test whether defective pyroptosis of these microglial subsets influences normal brain functions, we conducted a variety of behavioral assays on *Casp1*^*-/-*^ mice (Sup. Table 1)^17^. Despite previous research associating an inflammasome protein *NLRP1* with ASD^8^, we did not observe defective sociability (Sup. Fig. 2a), repetitive restricted behavior (Sup. Fig. 2b) or learning and memory deficits (Sup. Fig. 2c), which are common ASD-like behavior changes. Instead, we found that *Casp1*^*-/-*^ mice showed features of hyperactivity (Fig. 2a) and inattention (Fig. 2b) determined by the open field assay and 5-choice serial reaction time task assay, respectively. Interestingly, *Casp1*^*-/-*^ mice frequently climbed up the stranger mouse cylinders (with or without stranger mice) in the 3-chamber interaction assay and some mice eventually escaped from the test chamber (Fig. 2c and Sup. Movie 3 and 4). Inappropriate climbing behavior is considered to be a sign of impulsivity and often observed in children who are diagnosed with attention-deficit/hyperactivity disorder (ADHD)^18^. In addition, *Casp1*^*-/-*^ mice exhibited low levels of anxiety, as assessed by the elevated plus maze assay (Sup. Fig. 2d), which may contribute to the climbing behaviors and impulsivity of *Casp1*^*-/-*^ mice^19^. Of note, these behaviors changes were only seen in male offspring but not in females (data not shown), which is consistent with the male-bias of ADHD^20^. Interestingly, several reports suggest that the striatum and thalamus are constituents of neural circuits involved in hyperactivity and attention, respectively^21-23^. However, further studies are required to understand how defective microglial subsets of the LGE and iTh are involved in shaping these neuronal circuits.

**Fig. 2.**
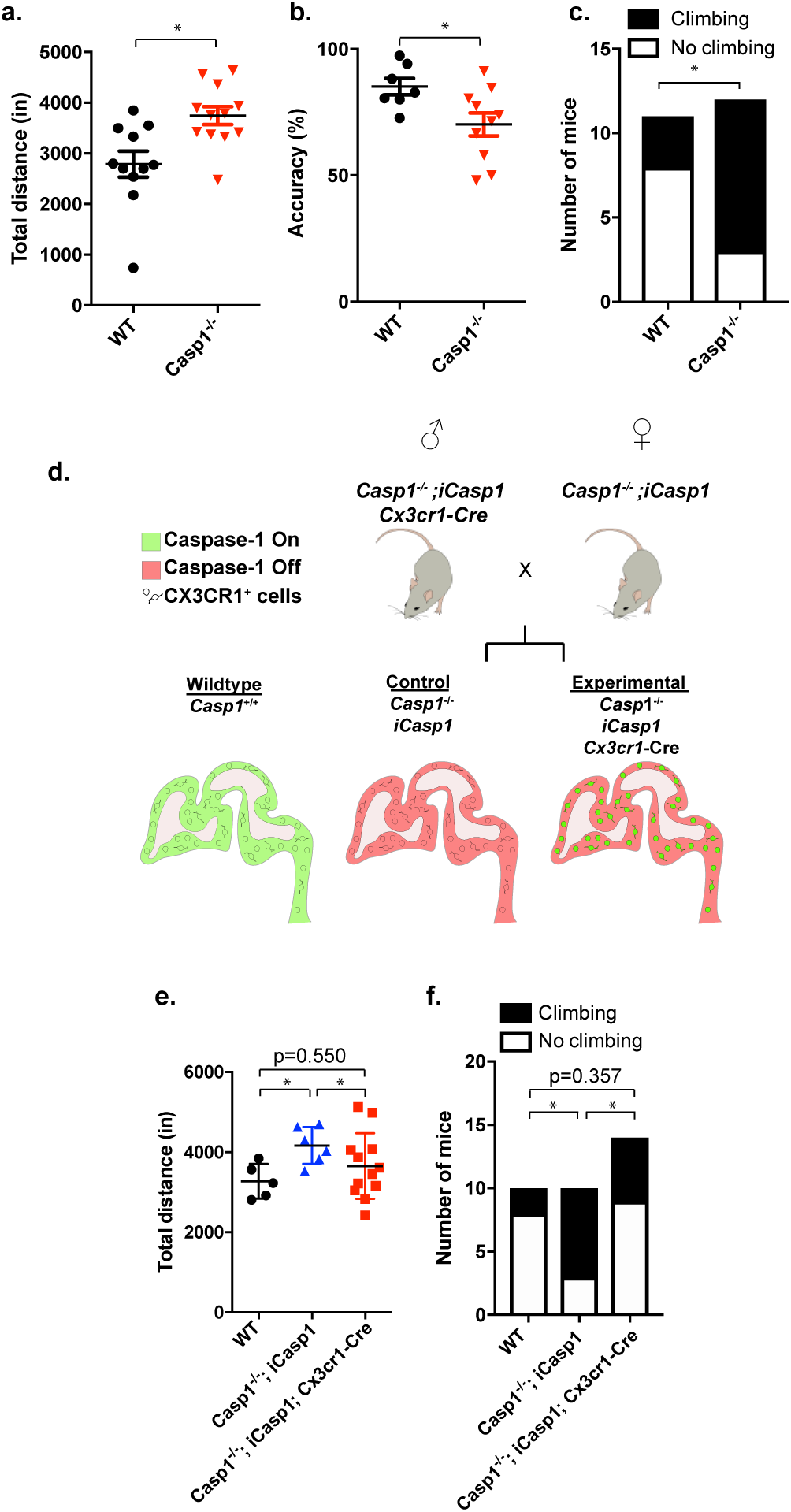
*Casp1* deficient mice exhibit ADHD-like behavior changes and Casp1 re-expression in Cx3cr1^+^ cells restores normal behavior. **a.** General activity of WT (black circles, N=11) and *Casp1*^*-/-*^ (red triangles, N=12) mice was determined by the open field assay and is shown as total distance traveled (inches). **b.** Attention behavior of WT (N=7) and *Casp1*^*-/-*^ (N=10) was determined by the five-choice serial reaction time task and is shown as accuracy of their response (%). **c.** Climbing behavior of WT (N=11) and *Casp1*^*-/-*^ (N=12) was determined by three-chamber social interaction assay. **d.** Breeding strategy is shown. We crossed three mouse lines (*Casp1*^*-/-*^, *iCasp1*, and *Cx3cr1*-Cre) to generate *Casp1*^*-/-*^; *iCasp1*; *Cx3cr1-Cre* positive mice (Experimental) and *Casp1*^*-/-*^; *iCasp1*; Cre negative littermates (Control). *Casp1* expression pattern depicted in sagittal embryonic brain illustrations. **e.** General movement of WT (black circles, N=5), littermate Control (blue triangles, N=6) and Experimental (red rectangles, N=12) mice was determined by the open field assay and is shown as total distance (inches). **f.** Climbing behavior of WT (N=10), littermate control (N=10), and Experimental (N=14) was determined by the three-chamber social interaction assay. Number indicates mice showed climbing behavior. * shows p<0.05.

The *Casp1*^*-/-*^ mice used in this study are conventional knockout mice in which *Casp1* is deleted from all cell types. We found that *Casp1* mRNA is highly expressed in Cx3cr1^+^ cells isolated from fetal brain cells, including microglia (Sup. Fig. 2e). In contrast, Cx3cr1^-^ cells displayed high expression of *Cd200* mRNA, which is associated with neural cells (Sup. Fig. 2f)^24^. In order to verify that expression of Casp1 in immune cells including microglia is necessary for ADHD-like behavior changes, we crossed *Casp1*^*-/-*^ mice to mice expressing a Cre-inducible *Casp1* allele (*Rosa26-LoxP-STOP-LoxP-Casp1-IRES-GFP,iCasp1* mice)^11^. These mice were then crossed to BAC transgenic mice expressing Cre under the control of the *Cx3cr1* promoter, a commonly used Cre-driver enriched in microglia^25,26^. The resulting *Casp1*^*-/-*^; *iCasp1*; *Cx3cr1*-Cre mice and their *Casp1*^*-/-*^; *iCasp1* littermate controls (equivalent to *Casp1*^*-/-*^ mice) were then used in our previously described behavior assays (Fig. 2d). Remarkably, re-expression of *Casp1* in Cx3cr1^+^ cells (Sup. Fig. 2g) restored normal behaviors (Fig. 2e-f and Sup. Fig. 2h). Upon Cre-mediated recombination, *iCasp1* mice also expressed GFP^11^, we found GFP^+^ cells that were positive for PI signal (Sup. Fig. 2i). These data suggest that *Casp1* deficiency is associated with behavior changes characterized by hyperactivity, inattention, and impulsivity and that re-expression of Casp1 in Cx3cr1^+^ cells including microglia restored the lytic cell death and normal behavior.

Since both GSDSMD and pro-IL-1β are cleaved by CASP1 (Sup. Fig. 1a), we also used *Gsdmd-* and IL-1 receptor (*Il-1r*)-deficient mice^11^ in our behavioral studies. These mice exhibited hyperactivity (Fig. 3a), and impulsivity (Fig. 3b and Sup. Fig. 3a), consistent with *Casp1*^*-/-*^ mice. In support with our observations, other reports described that mice deficient for IL-1 signaling exhibited ADHD-like behavior changes observed in our study^10,27,28^.

**Fig. 3.**
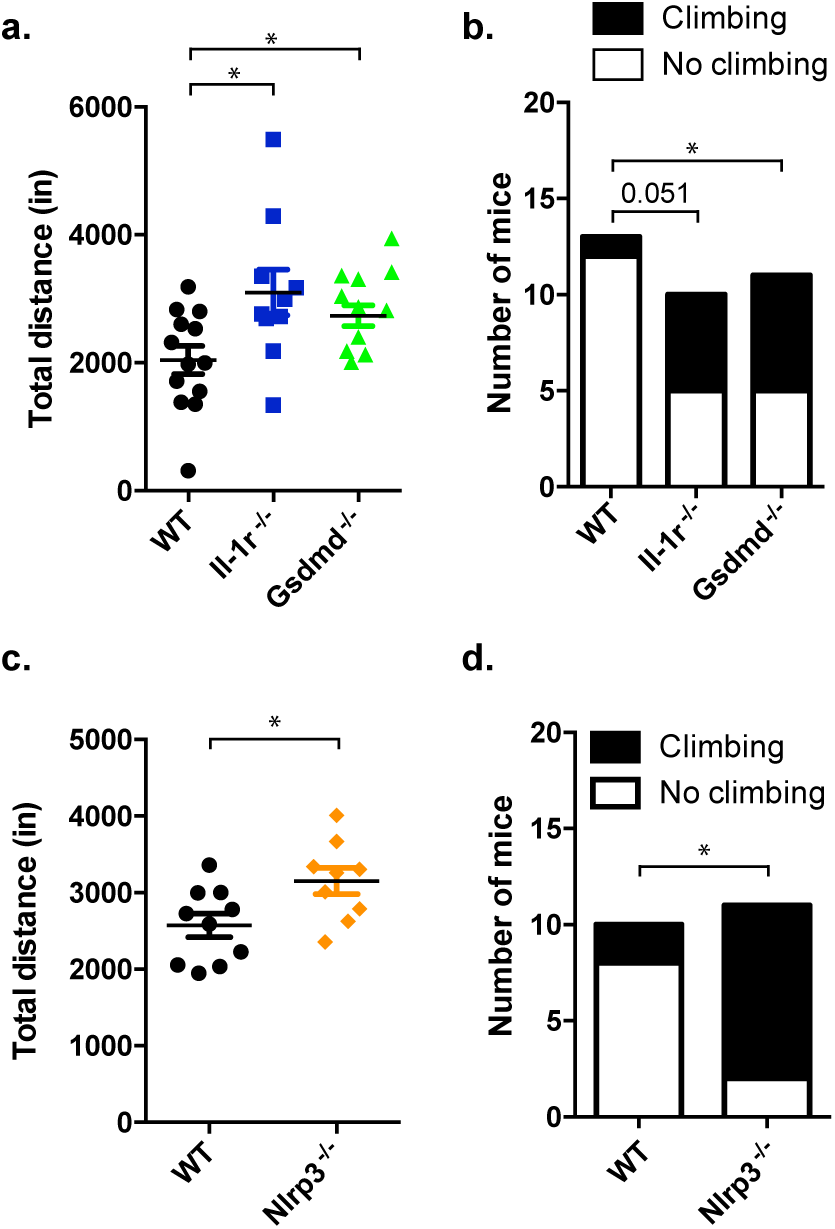
Mice deficient in the inflammasome/pyroptosis pathway exhibit ADHD-like behavior changes. **a.** General movement of WT (black circles, N=13), *Il-1r*^*-/-*^ (blue rectangles, N=10) and *Gsdmd*^*-/-*^ (green triangles, N=11) mice was determined by the open field assay and is shown as total distance (inches). **b.** Climbing behavior of WT (N=13), *Il-1r*^*-/-*^ (N=10), and *Gsdmd*^*-/-*^ (N=11) mice was determined by three-chamber social interaction assay. **c.** General movement of WT (black circles, N=10) and *Nlrp3*^*-/-*^ mice (orange diamonds, N=9) was determined by open field assay and is shown as total distance (inches). **d.** Climbing behavior of WT (N=10) and *Nlrp3*^*-/-*^ (N=11) mice was determined by three-chamber social interaction assay. Data are shown individual mice and error bars indicate S.E.M. Asterisk (*) shows p<0.05.

Next, we attempted to identify an upstream inflammasome molecule that was responsible for the initiation of the pyroptosis pathway (Sup. Fig. 1a). NLRP3 was a likely candidate because it can be activated by ATP and ions released from dying/dead cells. Since more than 50% of newly differentiated neurons die during fetal brain development^15^, these may activate NLRP3 and induce sterile inflammation during brain development (Sup. Fig. 1a)^29^.

To test this hypothesis, we ran our behavioral assays on *Nlrp3*^*-/-*^ mice and noted that they exhibited similar behaviors to *Casp1*^*-/-*^, *Il-1r*^*-/-*^, and *Gsdmd*^*-/-*^ mice (Fig. 3c-d, and Sup. Fig. 3b). From these results, we conclude that the activation of inflammasome that mediated by the NLRP3-CASP1-GSDMD/IL-1β pathway is required for normal mouse behavior and that loss of this pathway leads to ADHD-like behaviors.

Since our genetic experiments suggested that activation of the NLRP3-CASP1-GSDMD/IL-1R pathway in microglial subsets in LGE and iTh was important to prevent behavior changes, we tested if treatment of pregnant mice with specific drug inhibitors of CASP1 and NLRP3 (VX-765 and MCC950, respectively)^30^ may be sufficient to induce ADHD-like behaviors in their offspring. We injected WT pregnant mothers with CASP1 or NLRP3 inhibitors at E12.5 through E14.5, times when we observed fetal pyroptosis, and ran behavior assays on their offspring (Sup. Fig. 4). We found that male offspring from inhibitor-injected mice showed hyperactivity (Fig. 4a), and impulsivity, although to a lesser extent (Fig. 4b-c) when compared to our genetic models (Fig. 2 and 3). These data suggest that pharmacological disruptions of the NLRP3/CASP1 pathway, especially an inhibitor for NLRP3 (MCC950), may be sufficient to increase the risk of developing ADHD.

**Fig. 4.**
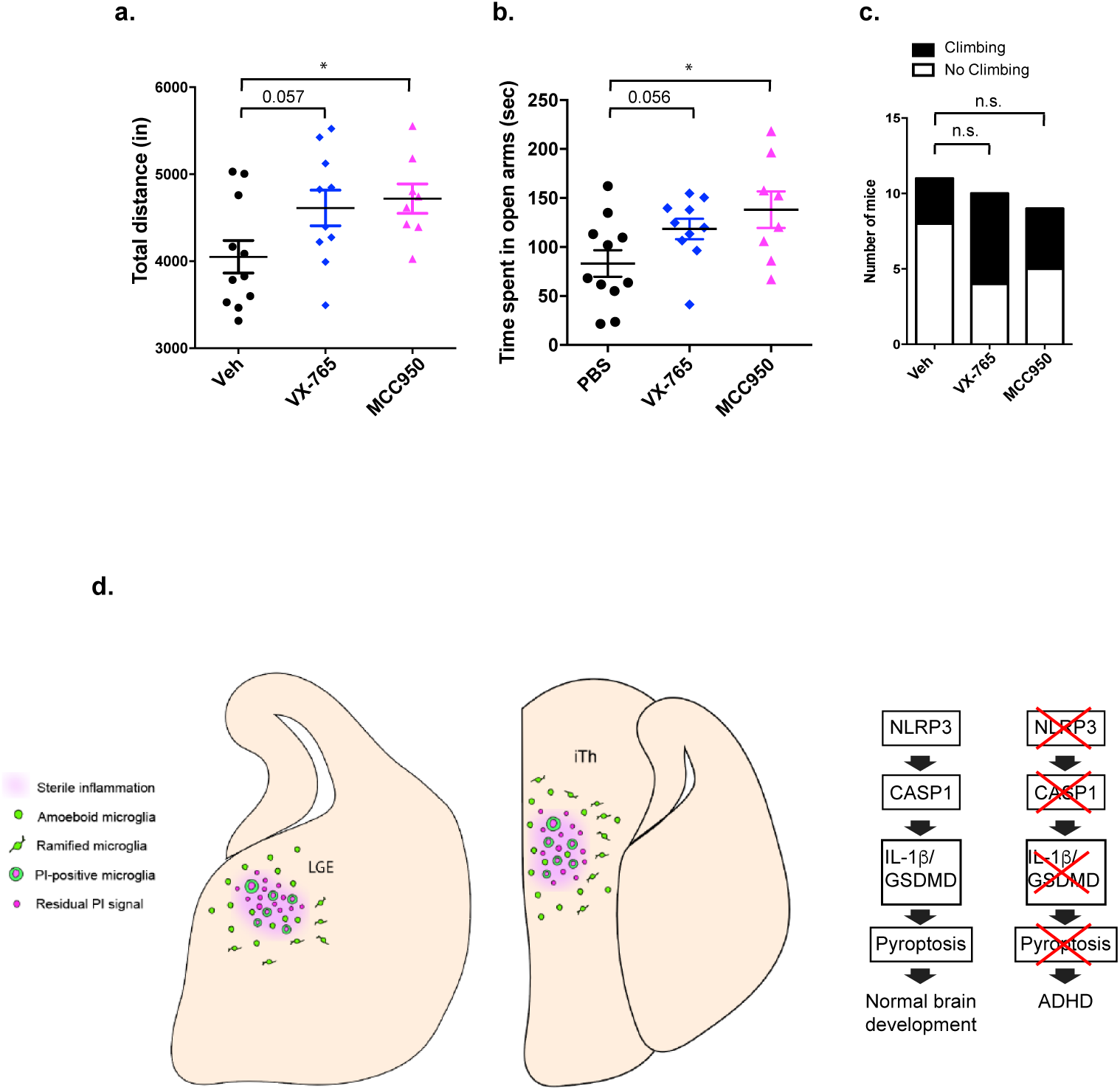
Fetal exposure to pyroptosis pathway inhibitor may lead to ADHD-like behaviors in mice. **a.** General movement of offspring of vehicle (Veh)-(black circles, N=11), VX-765,- (blue diamonds, N=10), or MCC950- (pink triangles, N=8) injected mothers was determined by open field assay and is shown as total distance (inches) traveled for each mouse. **b.** Level of anxiety of offspring of Veh- (N=11), VX-765- (N=10), or MCC950- (N=8) injected mothers was determined by the elevated plus maze assay and is shown as time spent in open arms (seconds). **c.** Climbing behavior of offspring of vehicle (Veh)- (N=11), VX-765- (N=10), or MCC950- (N=8) injected mothers was determined using the three-chamber social interaction assay. Data is shown as individual mice and error bars indicated S.E.M. Asterisk (*) shows p<0.05. **d.** Working model is shown. Specific microglial subsets in the LGE and iTh are characterized by pyroptosis during fetal brain development. Activation of the NLRP3-CASP1-IL-1β/GSDMD pathway is required for normal brain and genetic and pharmacological disruptions may increase risks of ADHD in offspring.

Overall, we have identified distinct subsets of microglia in LGE and iTh during fetal brain development which are characterized by the activation of inflammasome and pyroptosis pathways (working model, Fig. 4d). Furthermore, both genetic and pharmacologic alterations of this pathway result in ADHD-like behaviors in mice. Future work must investigate how dysregulation of these subsets of microglia are involved in shaping neuronal circuits underlying ADHD-like behaviors. Since microglia are highly heterogenous, further studies are required to understand how other microglial subsets may guide brain development and function.

## Material and methods

### Animals

All animals were maintained in specific pathogen-free conditions under a 12-hr light-dark cycle (7 am to 7 pm) and were given a standard chow diet and water *ad libitum*. Wild-type C57BL/6 mice were purchased from Charles River and The Jackson Laboratory. *Casp1*^*-/-*^*Casp11*^*-/-*^ mice were kindly provided by A. Van der Velden and M. Starnbach. The generation of *Casp1*^*-/-*^, *iCasp1* and *Gsdmd*^*-/-*^ mice was described previously^1^. Tg(Cx3cr1-cre)MW126Gsat mice were generated by Dr. Nathaniel Heintz at the Rockefeller University and purchased from MMRRC (UC Davis). *Il-1r*^*-/-*^ and *Cx3cr1*^*GFP/GFP*^ mice were purchased from The Jackson Laboratory. To normalized gut microbiota, all animals were cohoused in mixed-genotype groups of 3-5 mice per cage upon weaning. If cohousing was not done, their bedding was mixed regularly to normalize the microenvironment. All experiments were approved by the Animal Care and Use Committee and were performed under the institutional guidelines of the University of California, Berkeley.

### PI injection and tissue clearing

A total volume of 100 μl of propidium iodide (PI, 1.0 mg/ml) solution was intravenously injected into pregnant female mice at specific gestational dates. After 10-30 min of incubation, the mice were sacrificed. After recovering intact fetal bodies, we fixed fetal bodies and used the CUBIC tissue clearing method^2^. Briefly, embryos were incubated in R1 for approximately 2 days, replacing with fresh R1 after 24 hours, until the tissue was clarified (by eye). Embryos were rinsed in PBS before immersion in R2. They remained in R2, replacing with fresh R2 after 24 hours, until the tissue became fully clarified by eye. This step normally took 3-4 days before imaging.

### Light sheet fluorescence microscopy and analysis

LSFM was performed using the ZEISS Lightsheet Z.1 system and 5X objective set (EC Plan-Neofluar 5x/0.16 detection and LSFM 5x/0.1 illumination lens) at 0.36x zoom. Samples were mounted on the stage using superglue and FEP tubing. CUBIC R2 solution was used as the refractive index-matched solution. Coronal acquisitions captured the entirety of the embryonic forebrain (about 620 Z slices). GFP and PI signals were simultaneously acquired using one image track and 488 laser at 12% power, with two bandpass filters, 505-545 nm and 575-615 nm, to capture GFP and PI signals, respectively. Light sheet thickness set at 13.13 μm and exposure to 39.9 ms. Shannon-Nyquist sampling was observed, with voxel sizes approx. X =2.13 μm, Y=2.13 μm, and Z=6.16 μm. Files were converted to .ims format via Imaris File Converter and imported to Imaris 9.1.0 (Bitplane) for analysis. An embryo was counted as “cluster positive” if at least one cluster was found in the LGE or iTh. A “cluster” is defined as a group of at least 7 GFP^+^PI^+^ cells that formed a minimum surface volume of 7×10^5^ um^3^ containing residual PI signal. LGE and iTh regions were identified using anatomical features (such as proximity to ventricles and distance from most anterior plane) and comparing to standard embryonic mouse brain atlases (epmba.org and http://developingmouse.brain-map.org/). Further details are available upon request.

### Behavior assays

Male and female animals that were 2-4 months old were used for behavior assays.

#### Open field assay

Mice were individually placed in the center of a plastic box (22 × 42.5 × 21 cm) for 1 hour and allowed to freely explore the arena. Animal movement was monitored by computerized photobeam using the MotorMonitor SmartFrame System (Kinder Scientific).

#### Elevated plus maze assay

Mice were individually placed onto the center of a platform (5 × 5 cm) of the maze that consisted of two open and two closed arms (30 × 5 cm) that were elevated 30 cm from the floor. Mice were allowed to freely explore the maze for 10 min and the Smart Video Tracking System (Panlab) was used to determine the time spent in the open and closed arms, as well as the center, during the test.

#### Three-chamber interaction assay

A three-chambered rectangular box with two dividing walls that marked the left, center, and right chambers (20 × 40 × 22 cm, each) was used. Two walls had openings that allowed a test mouse to freely move into each chamber. After a 10-minute habituation phase, a cylindrical grid cage (7 cm diameter and 15 cm height) housing a stranger mouse (age- and sex-matched) (S1) and another empty cylindrical grid cage (E) were placed on opposite sides of the apparatus. The test mouse was allowed to freely explore the chambers with the S1 and E cages for 10 min. The movement of the test mouse was recorded, and the time spent interacting or sniffing each cylindrical cage was measured using the Smart Video Tracking System (Panlab). The time spent on the top of the cylindrical cage was designated as the climbing behavior and was also measured using the Smart Video Tracking System (Panlab) or analyzed manually.

#### Five-choice serial reaction time (5-CSRTT) assay

The procedures used in this task were as previously described with modifications^3^. A Plexiglas operator chamber (19 × 22 × 24 cm) with five response apertures and a food magazine was automatically controlled by the software (Packwin, Panlab). The procedure consisted of the pre-training, magazine training and 5-CSRTT training phase. Before the test and then throughout the entire experiment, the mice were food-restricted (1.8-2 g per day). On the first day of food restriction, mice were introduced some reward pellets (TestDiet 14 mg sugar pellets) to familiarize the mice to their taste. During the pre-training phase involved habituation to the chamber, five reward pellets in the food magazine and one reward pellet in each of the five apertures were placed for each operant chamber, and magazine light and all five stimulus lights were remained illuminated for the duration of the session. The mice were individually placed in the chamber and allowed to freely explore for 10 minutes. The habituation session was repeated until all the pellets were consumed. On the day of magazine training, the mice were placed in chamber for a 4-min period of free exploration, followed by all five stimulus lights switched on throughout the remaining session. After a random nose-poke response was made in one of the five apertures, the mouse was given 1 pellet in the food magazine. Once mice earned 20 pellets, they commenced to the 5-CSRTT training which graduates through increasingly challenging stages (stage 1-6) on a schedule of progressively decreasing stimulus duration (SD) and increasing inter-trial intervals (ITI). The percentage of accuracy was calculated as [(number of correct trials/ (number of correct and incorrect trials)) x 100]. The number of days taken to achieve the criteria for stage 6 (accuracy > 75 %, correct trials> 20) was recorded to indicate the attention performance. After meeting the criteria of stage 6 for at least two consecutive days, mice were moved forward in the testing schedule in which the variable ITI (5, 8.75, and 12.5 seconds) was randomly presented with an SD of 1.25 s. The percent accuracy was also calculated in this testing.

### Statistical analysis

Data are shown as averages with error bars indicating S.E.M. Sample sizes were determined based on our previous experiences and published works. Statistical analysis was performed using the Prism 7 (GraphPad) software. We used the Student’s t-test to compare two groups. Fisher’s exact test was used to correlate climbing behavior changes with genotype, as well as the appearance of clusters and their correlations with time point and genotype. P-values <0.05 were considered to be statistically significant.

## Acknowledgements

We thank the Saijo lab for discussion and D. Araujo, E. Robey and D. Kaufer for critical reading of the manuscript. We thank H. Aaron, J.Y. Lee, and F. Ives at the CRL MIC for imaging analysis help, and A. Sullivan (Obrizus Communications) for editing. K.S. is supported by start-up funding from the University of California, Berkeley, a National Institutes of Health (NIH) grant (1R01HD092093), a Searle Scholarship, a Hellman Fellowship, and a Pew Scholarship. R.E.V. is an HHMI Investigator and is supported by NIH grants AI075039 and AI063302. E.K.N. was supported by the Chancellor’s Fellowship, Elizabeth Roboz Einstein Fellowship, and University of California Dissertation-Year Fellowship. I.R. is supported by the Austrian Science Fund (the Erwin Schrödinger Fellowship). M.K. is supported by MCB departmental grant 4T32GM07232-40.

## Author information

These authors contributed equally and listed in alphabetical order of their last name: Hsiu-Chun Chuang, Eva K. Nichols.

## Author contributions

H-C.C. and E.K.N. designed and performed experiments and analyzed data. W-C.C., P.M.L., R.M., and M.K. performed experiments. I.R. and R.E.V. provided critical mouse lines and advice.

K.S. conceived the project and wrote the manuscript with input from all authors.

