## Supplemental material for "Defective cell death of distinct microglial subsets contributes to ADHD-like behavior in mice"

**Supplemental information**

**Supplemental Table 1. Behavior assay used in this study**

| | Symptoms | Behavior Assays |
|---|---|---|
| **Core** | Sociability | Reciprocal social interaction assay<br>Three-chamber interaction assay<br>Food preference social transmission assay |
| | Repetitive restricted Behavior | Marble burying assay |
| **Associated** | Learning/Memory | T-maze assay<br>Fear conditioning assay (contextual and cued) |
| | Anxiety | Elevated plus maze<br>Open field assay* (center vs peripheral ratio) |
| | Hyperactivity | Open field assay* (general and total activity) |
| | Attention | Five-choice serial attention task test |

* Using different parameters, two different behavior changes can be assessed by a single open field assay.

### Materials and Methods

#### Behavioral assay

T-maze assay

The T-maze assay was performed using an apparatus consisting of a start arm (15 x 5 x 12 cm) containing start area (8.9 x 5 x 12 cm) perpendicular to two opposing choice arms (15 x 5 x 12 cm, each) following the published protocol[1].

Marble burying assay

The procedures were described previously[2]. Briefly, the mouse was individually placed in a clean cage (30.5 x 16.5 x 14 cm) filled with the same bedding as the home cage (4.5 cm deep) for 20-min habituation period. Then, each mouse was returned to its home cage and 20 glass marbles (15 mm diameter) were gently overlaid in an equidistant 4 x 5 arrangement on top of the bedding. Mouse was allowed to access the marbles for 20 min and the number of marbles buried (>50% marble covered by the bedding materials) are recorded to test repetitive behaviors in mice.

**Dosing of VX-765 and MCC950**

The pregnant C57BL/6 mice were given three intraperitoneally injected doses of VX-765 (50 mg/kg/day, Caspase-1 inhibitor, InvivoGen), MCC950 (50 mg/kg/day, NLRP3-inflammasome inhibitor, InvivoGen), or 5% dimethyl sulfoxide (DMSO, control group), starting on embryonic day 12.5 until embryonic day 14.5 (Sup. Fig. 3).

**Immunofluorescence analysis**

Intact embryos were fixed in 4% PFA in PBS overnight at 4°C and underwent complete sucrose gradient up to 30% wt/vol in PBS. Embryos were next embedded in a 1:1 mix of O.C.T compound (Sakura VWR) with 30% sucrose in PBS in a -20°C chamber of a cryostat and sectioned at a thickness of 40 μm. Slides were dried and permeabilized with 0.1% Triton X-100 in PBS. After blocking, sections were incubated with Rb anti-hASC antibody (Adipogen, cat. no. AG-25B-0006; 1:200). After washing steps, sections were stained with Donkey anti-RbAF350 secondary antibody (ThermoFisher A10039; 1:1000). Sections were imaged using an upright Zeiss Axioskop 2 plus wide-field epifluorescence microscope. A 100 HAL lamp was used for illumination and images were captured with Plan Apochromat 20x/0.75 objective with standard DAPI and FITC cubes. An exposure time of 80.525 ms was used. Image brightness & contrast was changed to enhance visibility of ASC specks. At least five embryos were serial sectioned from three different litters to detect ASC signal in the forebrain. For *Casp1*$^{-/-}$*; iCasp1; Cx3cr1*-Cre staining, embryo was exposed to 100 μl of propidium iodide (PI) (i.v.) into the dam at least ten minutes prior to sacrifice. Chicken anti GFP primary antibody (Abcam ab13970; 1:1000) and goat anti chicken AF488 secondary antibody (ThermoFisher A11039; 1:1000, green) was used.

***Ex vivo* microglial isolation from fetal brain**

E14.5 fetal brains were individually dissected and minced in PBS, then placed in 5 mL of F-12 media containing 8 U/mL DNAse-I (Roche). Each sample was triturated, and samples were spun down and re-suspended in 10 mL of digestion buffer (HBSS buffer containing $Ca_2^+$ $Mg_2^+$ and 0.13 U/mL Liberase-TM, Sigma Aldrich) for 15 minutes. While preparing fetal brains, gDNA was isolated from fetal tails and used in a fast SRY PCR (Fwd 5'-TTGTCTAGAGAGCATGGAGGGC-3', Rev 5'-CCACTCCTCTGTGACACTTTAGC-3'; KAPA2G Fast HotStart PCR kit, Kapa Biosystems) to determine sex. Samples were pooled by gender and re-suspended in HBSS (without ions) containing 10% FBS. If fetal brains were Cx3cr1-$GFP^{+/-}$, samples were FACS sorted (FACS Aria II) as $GFP^+$ or $GFP^-$ cells. A viability dye (DAPI, 1:$10^4$) was used to detect dead cells. Samples were collected in Trizol-LS (ThermoFisher).

**RT-qPCR**

Total RNA from ex-vivo microglia was isolated using the DirectZol kit (Zymo research). RT reactions were performed using the SuperScript III First-Strand Synthesis System kit (Invitrogen, cat. no. 18080051) and qPCRs were performed using the KAPA SYBR qPCR 2X master mix (Kapa Biosystems, KK5503), all following the manufacturer protocols. $2^{-\Delta\Delta Ct}$ method was used for analysis using *Hprt* expression for normalization.  Primers used in this assay are following:

*Hprt* (Fwd 5'- TCAGTCAACGGGGGACATAAA-3', Rev 5'- GGGGCTGTACTGCTTAACCAG-3'), *Casp1* (Fwd 5'-TGGGACCCTCAAGTTTTGCCC-3', Rev 5'-GGCAAGACGTGTACGAGTGGTT-3'), and *Cd200* (Fwd 5'-GAAAGGCGCTGCACACAACTg-3', Rev 5'-GGCAGGCTGGATTACAACCCC-3').

**Supplemental Figure Legends**

**Sup. Fig. 1. Inflammasome activation pathway and PI$^+$ cluster characterization. a**. Activated inflammasomes serve as platforms to cleavage of pro-Caspase-1 to active Casp1, which then cleaves pro-interleukins (IL)-1β to generate mature cytokines. Casp1 also cleaves Gasdermin D (Gsdmd), which promotes subsequent pyroptosis, IL-1β secretion, and inflammation. **b.** A representative PI$^+$ cluster in the LGE at E12.5 as acquired by LSFM. Magenta: PI signal. **c.** Green: *Cx3cr*1-GFP. **d**. Merged. Scale bar indicates 200 μm. **e-h.** A representative visualization of the LGE region from an E14.5 fetal brain using **e.** bright field microscopy. **f.** Images within the rectangle are shown in *Cx3cr1*-GFP signal in grayscale. **g.** Anti-ASC staining signal in grayscale. Red triangles indicate specks. **h.** Merged image. Red triangles indicate specks. ASC signal is shown in magenta to enhance visualization.

**Sup. Fig. 2. *Casp1$^{-/-}$* male mice show low anxiety without autistic behavior changes, and *Casp1$^{-/-}$*; *iCasp1*; *Cx3cr1*-Cre mice characterization. a.** Sociability of WT (black circles, N=11) and *Casp1$^{-/-}$* mice (red triangles, N=12) was determined by the three-chamber interaction assay and the interaction time with a stranger mouse (S1) is shown (Δseconds). **b.** Repetitive restricted behavior of WT (N=11) and *Casp1$^{-/-}$* mice (N=12) was determined using the marble burying assay and the number of marbles buried (to 2/3 of their depth) is shown. **c.** Memory and learning of WT (N=6) and *Casp1$^{-/-}$* mice (N=6) was determined by the T-maze assay. Days required for 80% correct responses for three consecutive days are shown. **d.** Level of anxiety of WT (N=11) and *Casp1$^{-/-}$* (N=12) mice was determined by the elevated plus maze assay and is shown as time spent in open arms (seconds). **a-d**. Data is shown as individual mice and error bars indicate S.E.M. Asterisk (*) shows $p<0.05$. (ns) indicates not significant. **e-f.** *Casp1* (**e**) and *Cd200* (**f**) mRNA expressions in FACS-sorted *Cx3cr1*-GFP$^+$ (black bar) and *Cx3cr1*-GFP$^-$ (white bar) cells were isolated from E14.5 fetal brain and determined by RT-qPCR. Data are shown as average of duplicates and error indicated as S.E.M. Asterisks (****) indicate p<0.0001. **g.** *Cd11b*$^+$ cells were isolated from the brains of adult WT (black circles, N=3), littermate control (*Casp1*$^{-/-}$*; iCasp1*, blue rectangles, N=4), and Experimental (*Casp1*$^{-/-}$*; iCasp1; Cx3cr1*-Cre*,* red rectangles, N=4) mice and the mRNA expression of *Casp1* was determined by RT-qPCR. **h.** Level of anxiety of WT (black circles, N=7), littermate control (*Casp1*$^{-/-}$*; iCasp1*, blue rectangles, N=10), and Experimental (*Casp1*$^{-/-}$*; iCasp1; Cx3cr1*-Cre*,* red rectangles, N=12) were determined by elevated plus maze assay. Data is shown as individual mice and error bars indicate S.E.M. Asterisk (**) indicates p<0.01 and (****) indicates p<0.0001. **i.** A representative image of PI$^+$ cluster in the LGE at E14.5 of *Casp1*$^{-/-}$*; iCasp1; Cx3cr1*-Cre mice (N=2) is shown. Bright field (top left), GFP (top right), PI (bottom left), and merge (bottom right). Scale bar indicates 220 μm (bright field) and 130 μm (fluorescent staining). The contrast is increased in PI imaging for better visualization.

**Sup. Fig. 3. Mice deficient for Gasmd and Nlrp3 show low anxiety level. a.** Level of anxiety of WT (black circles, N=13), *Il-1r*$^{-/-}$ (blue rectangles, N=10) and *Gsdmd*$^{-/-}$ (green triangles, N=11) mice was determined by elevated plus maze assay and is shown as time spent in open arms (seconds). **b.** Level of anxiety of WT (N=10) and *Nlrp3*$^{-/-}$ (N=9) mice was determined by elevated plus maze assay and is shown as time spent in open arms (seconds). Data is shown as individual mice and error bars indicate S.E.M. Asterisk (*) shows p<0.05, and double-asterisk (**) shows p<0.01.

**Sup. Fig. 4. Pharmacological disruption of pyroptosis pathway may lead to ADHD-like behavior.** Experimental scheme is shown. VX-765 or MCC950 or vehicle as a control was injected to WT pregnant female at E12.5, E13.5, and E14.5. Behavior assays of offspring of compound injected mothers were performed when they were 10-weeks old.

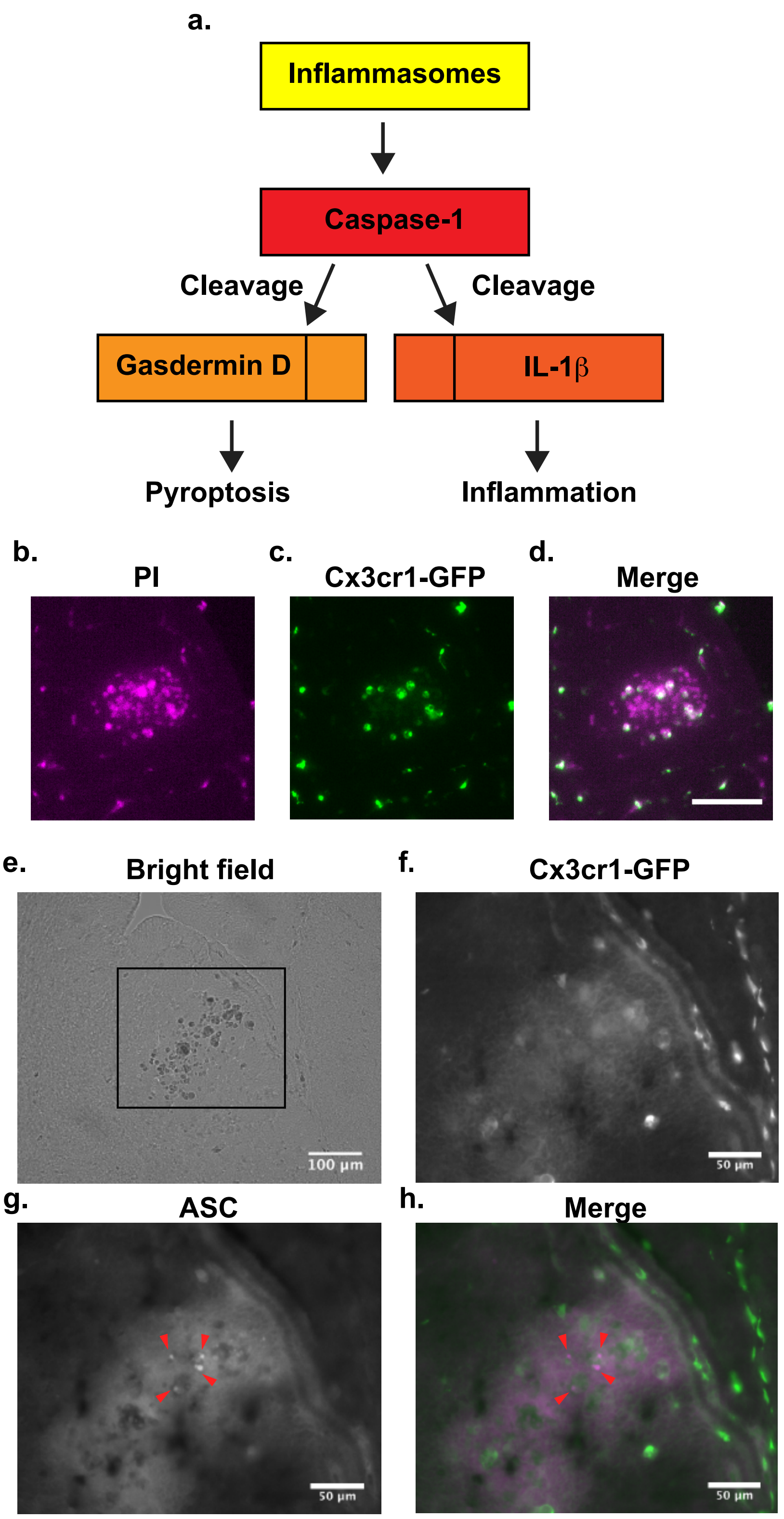


Chuang and Nichols et al. Sup. Fig. 1

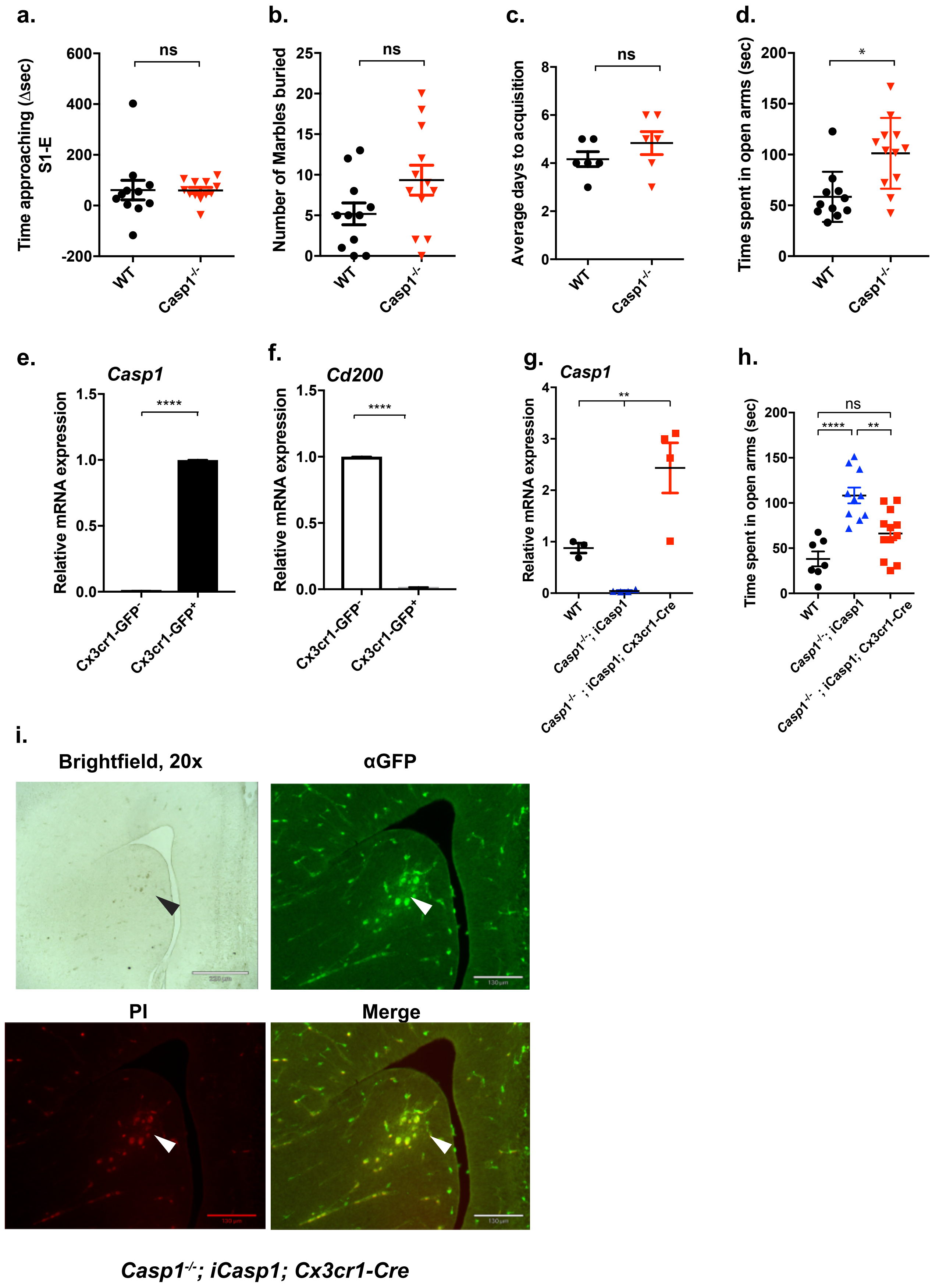

a.
b.
c.
d.
e.
f.
g.
h.
i.
Time approaching (Δsec) S1-E
Number of Marbles buried
Average days to acquisition
Time spent in open arms (sec)
Relative mRNA expression
Casp1
Cd200
ns
*
****
**
WT
Casp1-/-
Cx3cr1-GFP-
Cx3cr1-GFP+
Casp1-/-; iCasp1
Casp1-/- ; iCasp1; Cx3cr1-Cre
Brightfield, 20x
αGFP
PI
Merge
Casp1-/-; iCasp1; Cx3cr1-Cre

a

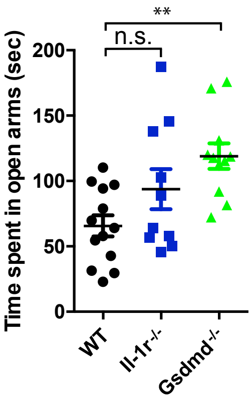

**
n.s.
Time spent in open arms (sec)
200
150
100
50
0
WT
Il-1r-/-
Gsdmd-/-


b.

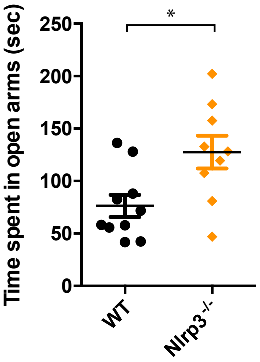

*
Time spent in open arms (sec)
250
200
150
100
50
0
WT
Nlrp3-/-

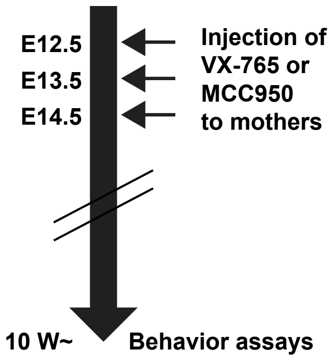


Chuang and Nichols et al. Sup. Fig. 4
